# A step toward synthetic endosymbiosis: introducing *Escherichia coli* and the trypanosomatid endosymbiont *Candidatus* Kinetoplastibacterium crithidii into mammalian cells using a polyethylene glycol-based fusion protocol

**DOI:** 10.64898/2026.09.20.753003

**Authors:** Rudvi Pednekar, Natascha A van Geelen-Kuenzel, Carroll MC Diehl, Leonie-Alexa Koch, Briardo LLorente, Eva CM Nowack, Matias D Zurbriggen

## Abstract

Synthetic endosymbiosis, the deliberate introduction of bacteria into eukaryotic host cells, provides a tractable model for how endosymbiotic relationships become established, and a route to cells with new metabolic capabilities that never evolved naturally in that lineage. Current methods for introducing bacteria into mammalian cells rely on microinjection, engineered invasion, listeriolysin-mediated phagosomal escape, or naturally permissive hosts. These methods are invasive, low-throughput, or costly, and most require the prospective endosymbiont to be genetically tractable, which excludes endosymbiotic bacteria which are of great interest as potential organelle precursors. Here we describe a protocol based on polyethylene glycol (PEG), long used to fuse mammalian cells, that delivers *Escherichia coli* into HeLa cells without genetic modification of either partner or the use of specialised equipment. We optimised the governing parameters in two stages: first, bacterial density, gentamicin concentration, and PEG molecular weight and concentration were optimised; then, medium composition and the timing of fusion and recovery. Across four parameters screened, 10% PEG-3350 applied for 4 min to HeLa cells 12 h after seeding, followed by 4 h in gentamicin-containing medium, gave the most consistent delivery. Under these conditions, confocal microscopy and z-stack reconstruction detected mCherry-tagged *E. coli* inside HeLa cells. Applying the same protocol to *Candidatus* Kinetoplastibacterium crithidii, a naturally occurring β-proteobacterial endosymbiont of the trypanosomatid *Angomonas deanei*, we observed the endosymbiont inside the HeLa cells. This protocol offers an accessible entry point for constructing artificial endosymbioses.

## Introduction

Endosymbiosis has repeatedly driven the evolution of eukaryotic cellular complexity. The transformation of endosymbiotic bacteria into genetically integrated organelles was central to the origin and diversification of eukaryotes and development of new metabolic and physiological capabilities. It gave rise to mitochondria and plastids and reshaped the trajectory of cellular complexity. This process starts with the integration of one organism into another, often leading to mutual benefits such as nutrient exchange or protection. The Endosymbiosis theory suggests that many bacterial genes in eukaryotic genomes originated from organelle ancestors that entered the eukaryotic lineage^1,2^. The transformation of a free-living bacterium into a genetically integrated organelle was gradual, and the intermediate stages are largely inaccessible, as the events that produced mitochondria and plastids occurred more than a billion years ago.

Merging two cells, or introducing one cell into another, represents an emerging frontier in synthetic biology. Synthetic endosymbiosis aims to reconstruct this process in the laboratory, both to gain experimental access to an ancient transition otherwise inferred from comparative genomics, and to build cells with metabolic capabilities which never evolved naturally with potential applications in biotechnology, medicine and evolutionary biology.

Synthetic endosymbiosis has most often been attempted in permissive hosts like yeast, in which auxotrophic *E.coli* has been sustained over many generations^3^. Mammalian cells are far more demanding hosts, but also a far more consequential one; an engineered endosymbiont could supply metabolic functions a human cell lacks or actively direct its behaviour as demonstrated by bacteria engineered to secrete mammalian transcription factors that redirect host cell fate^4^. To establish synthetic endosymbioses in mammalian cells, a few principal barriers have to be considered, such as phagolysosomal degradation, anti-growth cytoplasmic factors, antigen presentation via MHC pathways, and the induction of apoptosis or autolysis in response to the associated stress^5^.

Various technologies are implemented to introduce bacteria or organelles into eukaryotic host cells. Isolated spinach chloroplasts internalised by mouse fibroblasts degraded within approximately five cell generations^6^. Cyanobacteria engineered to express invasin and listeriolysin O persisted for approximately twelve hours in macrophages and induced autolysis in CHO cells^7^. The most durable associations reported to date include auxotrophic *E. coli* maintained in yeast for 40-120 generations^3^. Bacteria have been introduced into mammalian host cells by several routes. Microinjection has delivered *Listeria monocytogenes, Shigella flexneri* and enteroinvasive *E. coli* (EIEC)^8^ and *Salmonella enterica* serovar Typhimurium (*S.* Tm)^9^ directly into the cytosol, and fluid force microscopy^10^ has been used to inject *E.coli*^10^. Host cells have also been engineered to permit invasin or listeriolysin mediated invasion^7,10^; and ciliate *Tetrahymena thermophila* has served as a naturally permissive host^11^. These protocols are invasive, low-throughput, and expensive. For example, microinjection and fluidic force microscopy (FluidFM) operates on individual cells and requires dedicated instrumentation.

Some bacterial targets pose subsequent constraints, as exemplified with *Candidatus* Kinetoplastibacterium crithidii, a ß-proteobacterial endosymbiont of the trypanosomatid *Angomonas deanei*. The endosymbiont possesses a reduced genome, divides in synchrony with its host, is not genetically tractable and has not been yet cultured axenically^12–15^. Several host-encoded proteins are targeted to this endosymbiont, indicating a degree of host control over endosymbiont integration^14,16^. It therefore represents an advanced endosymbiosis and an informative system in which to study host-endosymbiont integration. However at the same time it is inaccessible to every delivery method mentioned above as every established route into a mammalian cell requires the bacterium to participate: to be transformed with invasion or escape factors, to survive culture beforehand, or to be introduced individually by micromanipulation. A bacterium that cannot be cultured, cannot be modified, and remains viable for only a few hours, meets none of these conditions. Instead, we need a method in which the bacterium is inert, the medium mediates crossing of the host cytoplasmic membrane, and the bacterium thus only needs to be present.

Polyethylene glycol (PEG)-based fusion would be such a method. PEG is a neutral, highly hydrated polymer that binds substantial quantities of water. When two membranes are apposed in concentrated PEG, the polymer withdraws water from the intervening space, the resulting dehydration is thought to produce an asymmetry in lipid packing pressure between the two leaflets of each bilayer, forming a single bilayer septum at the point of contact that then decays as the initial fusion pore opens^17^. As the driving force is physical rather than receptor-mediated, no recognition system, invasion apparatus, or genetic modification is required of either partner. Both, molecular weight and concentration of the PEG, influence the outcome, since together they determine the extent of water withdrawal and viscosity of the medium.

PEG has been used to induce fusion between plant protoplasts^18^ and between yeast protoplasts and hen erythrocytes^19^, and Pontecorvo et al. (1971) demonstrated PEG-mediated fusion of mammalian cells with subsequent recovery of viable hybrids^20^. The absence of surface specificity has permitted fusions that no biological mechanism would support; HeLa cells were fused with tobacco protoplasts in 1976, yielding viable interkingdom hybrids that survived six days^21^, and human and *Arabidopsis* cells have more recently been fused to generate proliferating lines carrying chromosomes from both partners, in which plant genes were transcribed by the human machinery^22^. Cell fusion has been identified as one of three principal routes to artificial photosynthetic animal cells^23^, and interkingdom fusion has been proposed, though not tested, as a strategy for artificial endosymbiosis^5^.

A PEG-based protocol would therefore differ from existing methods in three respects: (i) the bacterium is a passive participant, so neither culture nor genetic modification is required of it, (ii) the procedure acts on whole populations in a single step, rather than one cell at a time and, (iii) it needs no complex instrumentation beyond a pipette. Together these traits place organisms such as *Ca*. K. crithidii within reach to establish a synthetic endosymbiosis for the first time.

To our knowledge, PEG has not previously been used to deliver bacteria into mammalian cells. Two problems must be addressed for this to be feasible. First, the property that enables fusion might also limit it: hyperosmotic conditions slow *E. coli* growth by causing water loss and loss of turgor in cells. PEG at the concentrations used for mammalian cell fusion could therefore harm the bacteria^24,25^. Since this effect of PEG has not been well characterised, we first identified the range of PEG molecular weights and concentrations that do not impair bacterial growth. Second, bacteria proliferate rapidly in mammalian culture medium, and extracellular growth must therefore be suppressed without compromising bacteria already delivered or the viability of the host.

Here, we address both possible limitations using *E. coli* as a prototyping chassisOptimisation, requires bacteria in bulk and on repeated occasions, since each parameter must be tested across a range of values. *E. coli,* which can be cultured to any required density on demand. Bacterial density and gentamicin concentration, PEG molecular weight and concentration, medium composition and incubation timings were systematically optimised to yield conditions that reproducibly deliver *E. coli* into HeLa cells. The resulting protocol was then successfully applied to the endosymbiont *Ca*. K. crithidii isolated from *A. deanei,* indicating its potential as a general route for constructing artificial endosymbioses. Isolation of *Ca.* K. crithidii requires successive density gradient centrifugation steps, yields limited material, and produces endosymbionts that remain viable for only a few hours. Therefore, as a general approach setting up the protocol in E. coli first, which is comparable in size to *Ca*. K. crithidii making it a suitable proxy, and afterwards in the target endosymbiont is a straightforward approach.

### Protocol development and optimisation for endosymbiont - mammalian host cell PEG fusion

The procedure must balance two sets of competing constraints: it has to be mild enough so that bacteria and host cells survive, but intense enough to enable fusion. We developed and optimised the method using *E. coli* as a prototyping chassis to screen the full parameter space at each stage while carrying forward only conditions that satisfied both the bacterium and host cell, and afterwards implemented it to fuse *Ca.* K. crithidii with the mammalian cell (Fig. 1).

**Figure 1.**
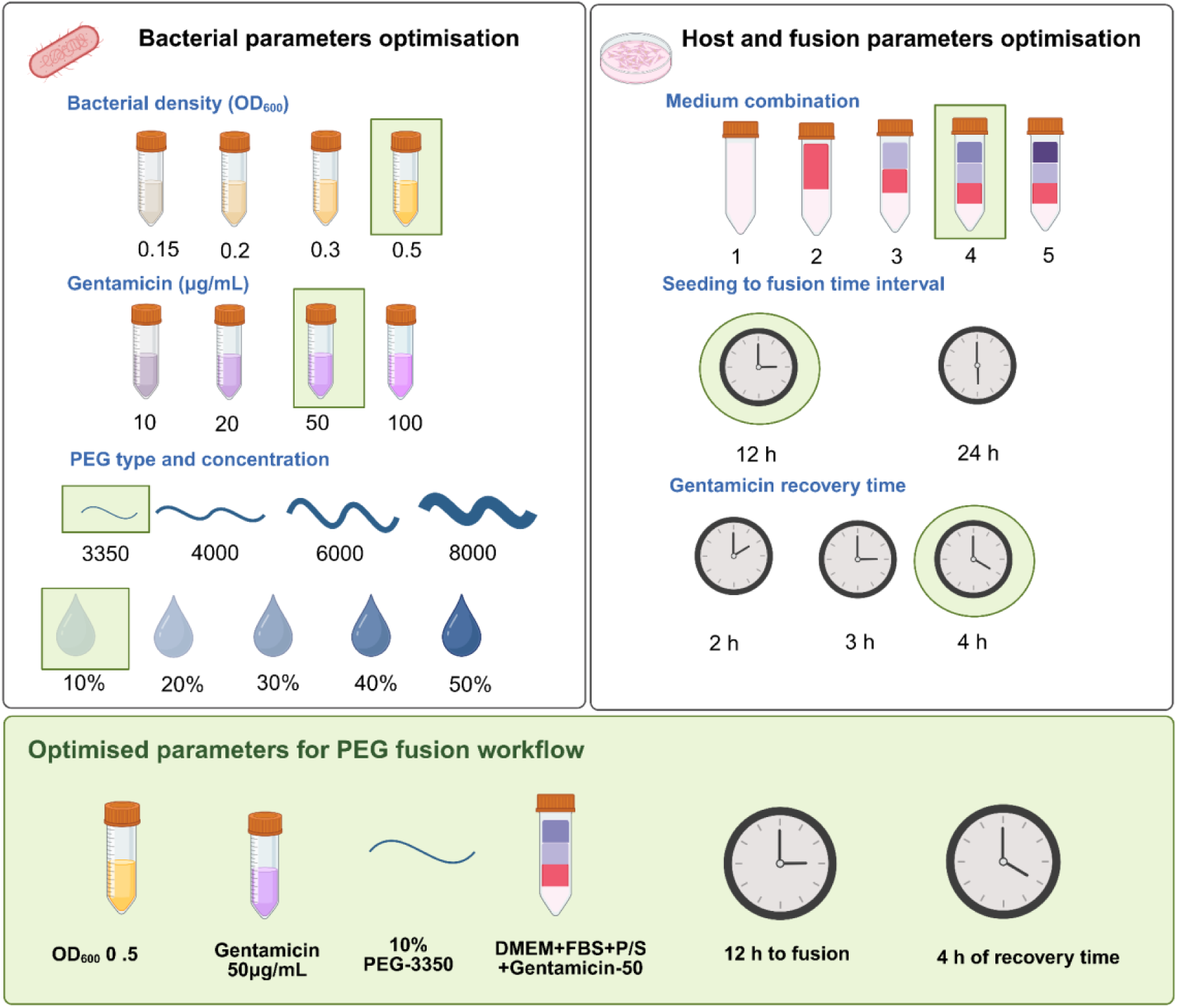
Two-part optimisation strategy. Optimisation was carried out in two stages. Bacterial parameters optimisation established the constraints set by the bacterium, and comprised screening gentamicin concentration against bacterial density (16 conditions) and PEG molecular weight against PEG concentrations (20 conditions). This fixed the bacterial optimal input OD_600_ 0.5, gentamicin at 50 µg mL^-1^and PEG-3350 at 10-20%. Host and fusion parameters optimised within those constraints, screening medium composition (5 combinations; 1: DMEM, 2: DMEM+FBS 3: DMEM+FBS+P/S 4: DMEM+FBS+P/S+gentamicin 50 µg mL^-1^ 5: DMEM+FBS+P/S+gentamicin 100 µg mL^-1^) and the timing of PEG exposure, seeding and recovery. Highlighted box showing the optimised parameters for the PEG fusion workflow.

#### Optimisation of bacterial parameters

The bacterium establishes its boundary conditions considering that two steps in the process are actively detrimental to its viability: i) the antibiotic of choice that should inhibit growth and proliferation of extracellular bacteria, without affecting the viability of the intracellular ones, and ii) the fusion reagent used. As previously employed in Gäbelein et al.^10^ we used gentamicin to eliminate the extracellular bacteria.

##### Bacterial density and gentamicin concentration

Since bacteria can proliferate exponentially in the rich mammalian cell culture medium, we aimed to limit proliferation exclusively to bacteria within HeLa cells. To address this issue, we considered the antibioticum gentamicin, which specifically targets the 30S subunit of bacterial ribosomes, disrupting translation^26^. As a polycationic aminoglycoside, it crosses the mammalian plasma membrane poorly, so intracellular bacteria remain unexposed^26,27^, whereas growth of bacteria located outside mammalian cells is effectively suppressed. We evaluated gentamicin concentrations of 10 µg, 25, 50, and 100 µg mL^-1^ against the bacterium at four bacterial densities (OD_600_ 0.15, 0.2, 0.3, and 0.5), yielding 16 combinations (Suppl. Fig. S1). All concentrations tested suppressed growth with OD_600_ peaking at approximately 2 h and suppression beginning thereafter, whereas the antibiotic free controls continued to increase. We selected 50 µg mL^-1^ with the bacterial input OD_600_ of 0.5 (Fig. 2A-B), considering that higher bacterial concentration than 0.5 OD might be harmful for HeLa cells. Gentamicin is reported to be non-toxic to mammalian cells at these concentrations^27^ and we observed no effect on HeLa morphology or attachment; therefore, additional tests of gentamicin concentration effects on HeLa cell viability were not conducted.

**Figure 2.**
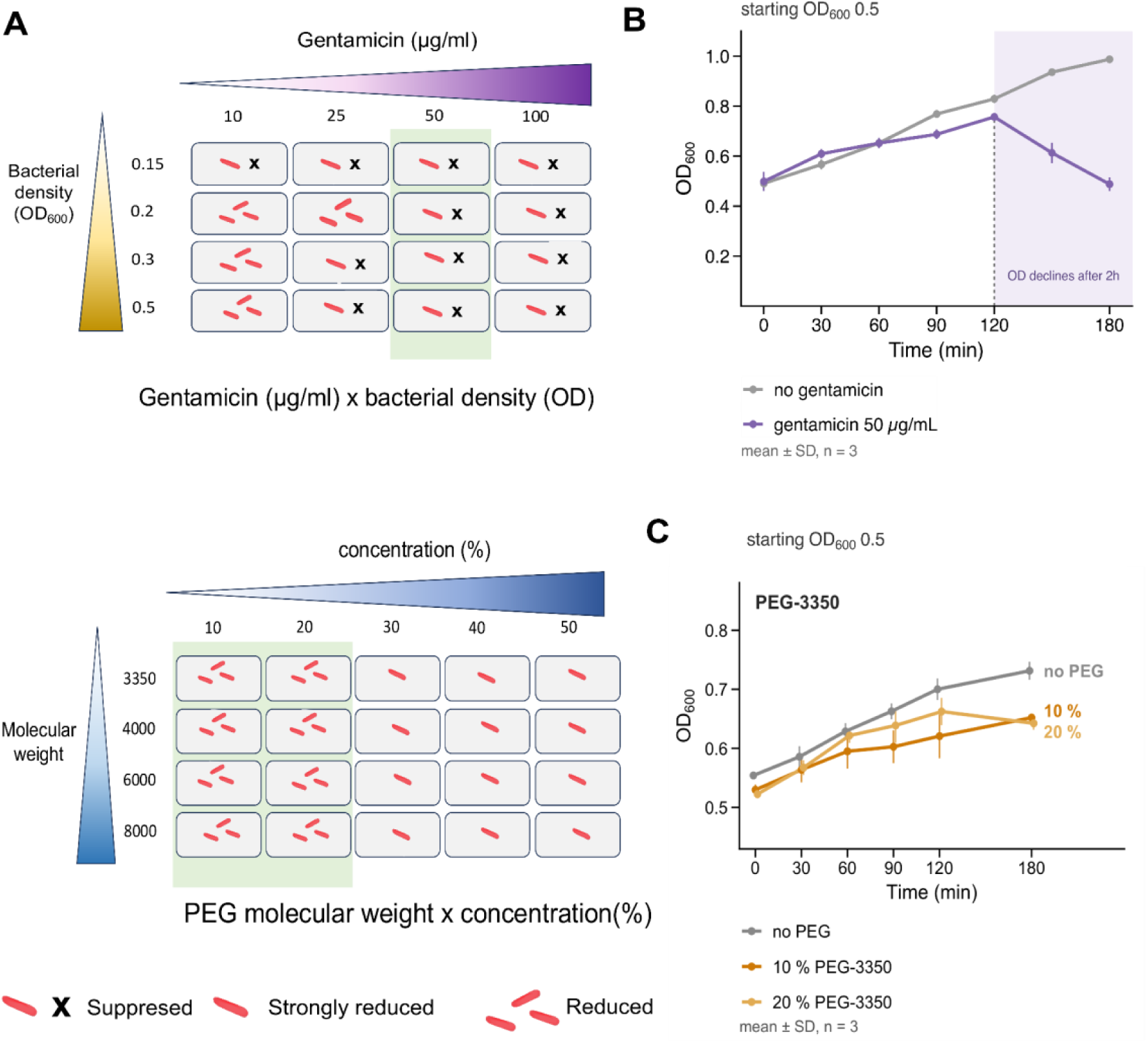
Optimisation of bacterial parameters. **(A)** Screened parameters: gentamicin (10, 25, 50, 100 µg mL^-1^) was tested with bacterial density (OD_600_ 0.15, 0.2, 0.3, 0.5), and PEG molecular weight (-3350, -4000, -6000, -8000) at concentrations 10-50%. The highlighted parameters were selected for the next experiments. Complete matrices are given in Suppl. Figs. S1, S2. **(B)** Growth of *E. coli* at the selected gentamicin concentration across the 4 densities was tested with an antibiotic-free control; growth suppression began at approximately 2 h. **(C)** Growth at OD_600_ 0.5 in 10% and 20% PEG-3350 with a no-PEG control. Values are shown as a mean 土 SD of three technical replicates from a single experiment.

##### Polyethylene glycol weight and concentration

As a fusion reagent, we selected PEG, which is widely used in cellular fusion procedures^20,28,29^. We then aimed to identify the maximum PEG concentration compatible with bacterial viability. Fusion efficiency between mammalian cells increases with PEG concentration, with little fusion reported below 30% w/w^29^. These thresholds were established for fusion between two mammalian cells, and do not necessarily need to apply to the delivery of a bacterium, which differs in size, envelope composition and contact geometry. However, as mentioned before, PEG could restrict bacterial growth. Therefore, we combinatorially tested multiple PEG formulations at different concentrations to find the optimal conditions. Four molecular weight PEGs (PEG-3350, PEG-4000, PEG-6000, and PEG-8000) at five concentrations (10%, 20%, 30%, 40%, and 50%) were tested against a bacterial culture with an OD_600_ of 0.5 (Fig. 2A; Suppl. Fig. S2). Growth was reduced relative to the no-PEG control at every concentration tested and the reduction increased with the concentration. Molecular weight had little systematic effect. Growth in 10% and 20% PEG-3350 remained comparable to the no-PEG control for approximately 90 min, after which the OD slightly reduced for both the PEG-3350 and PEG-4000 compared to no-PEG control (Fig. 2C).

Based on the results, we used a PEG concentration of 10-20% for subsequent experiments, since PEG at these levels is not toxic to mammalian cells^30^.

In sum, the optimisation of bacterial parameters established three parameters for the resulting PEG-fusion protocol: a bacterial concentration with OD_600_ of 0.5, a gentamicin concentration of 50 µg mL^−1^ (Fig. 2B) and 10% to 20% PEG-3350 (Fig. 2C). Out of these conditions, OD_600_ of 0.5 and a gentamicin concentration of 50 µg mL^−1^ were maintained throughout the subsequent optimisation process and 10% and 20% of PEG-3350, -4000, -6000 and -8000 were further tested in fusion tests.

#### Optimisation of Host and PEG fusion parameters

As mammalian cell host, we selected HeLa cells based on metabolomics evidence indicating that the predominant portion of their metabolite pool comprises free amino acids, accounting for approximately 80% of the metabolites associated with central carbon metabolism. These amino acids function as a significant source of carbon and energy, thereby eventually supporting the proliferation of cytosolic *E. coli*^10,31^.

Having the bacterial (co)-culture conditions fixed, we optimised the parameters for the host cell and fusion steps itself: These are governed by a different and partly opposing set of requirements. The culture medium must sustain bacterial viability sufficiently for the fusion to occur, albeit also facilitating the suppression of residual extracellular bacteria while supporting host cell viability. This presents a challenge, as a medium that inhibits bacterial growth initially would hinder the delivery, whereas one that permits continuous growth could lead to overculturing. The interval between seeding of the mammalian cells and fusion influences the host monolayer morphology; a short interval may not allow adequate cell spreading, reducing available surface area after fusion; conversely, a longer interval may lead to monolayer confluence potentially affecting the delivery of bacteria inside the host cell. The duration of PEG exposure must balance delivery efficiency against host cell viability, as dehydration enhances membrane contact but stresses the cells thus reducing viability. Following fusion, the recovery time must be optimised to ensure effective extracellular bacterial clearance by gentamicin without compromising host integrity. Hence, three parameters were optimised: medium composition, seeding-to-fusion interval, and recovery time post-fusion in gentamicin. Unlike bacterial growth assays, these parameters were assessed primarily by delivery efficacy rather than bacterial viability alone. The effect of medium composition was evaluated with growth curves (Fig. 3A-B), clearance dynamics by taking confocal images at different time intervals (Fig. 3C), and overall delivery success was determined by the presence of mCherry-positive *E. coli* cells inside of HeLa cells. The timing matrix (Fig. 3D) shows the various timepoints and conditions tested.

**Figure 3.**
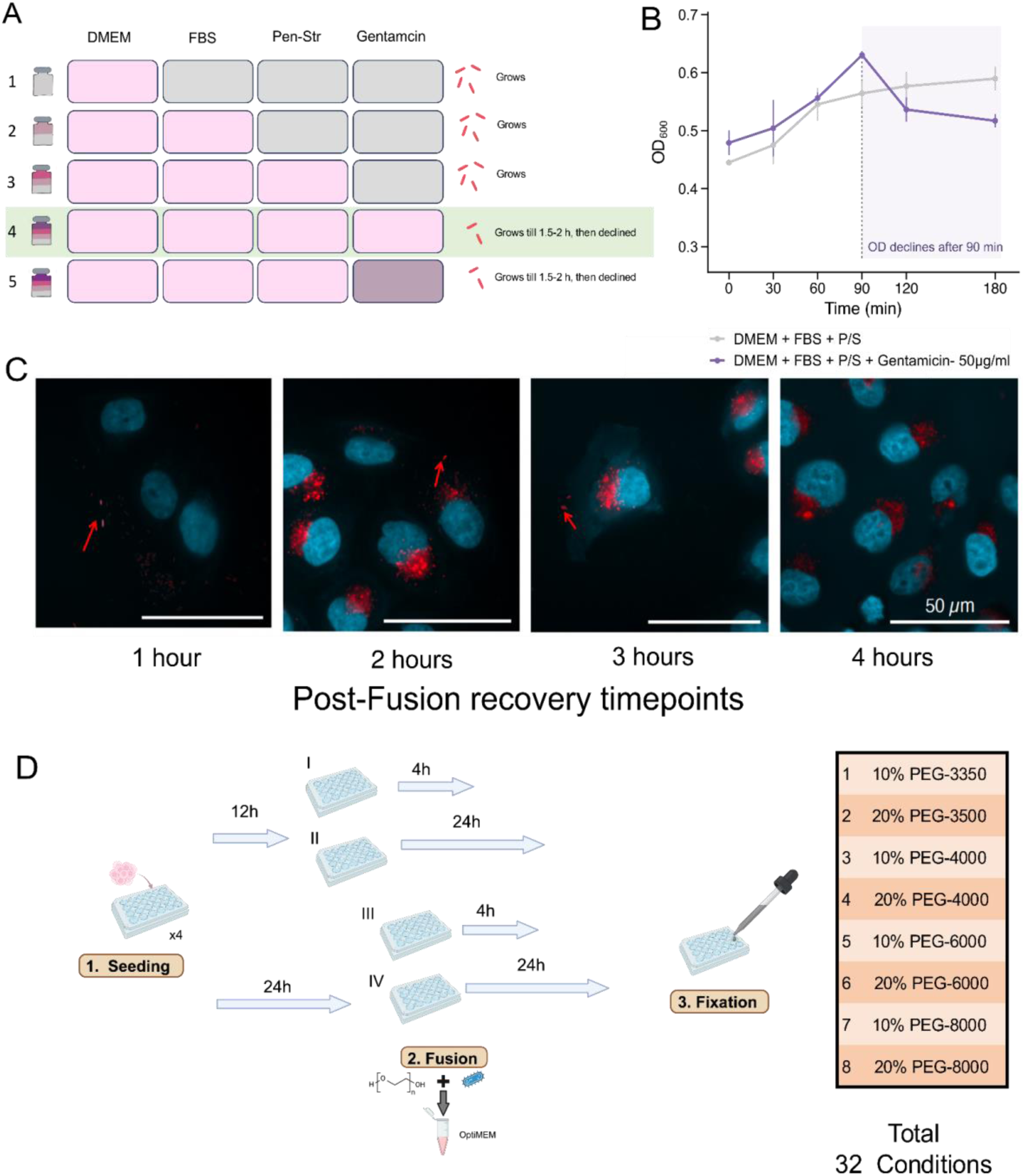
Optimisation of host and fusion parameters. **(A)** Five cumulative DMEM-based medium combinations were screened. Bacterial growth was unaffected by FBS and bacterial growth was cleared faster in the presence of gentamicin than in the medium with only penicillin and streptomycin. Combinations 4 and 5 both met the requirement, and combination 4 was used in the final protocol on the basis of the concentration fixed in the bacterial parameter optimisation. **(B)** Growth of *E. coli* in combination 4 compared with a gentamicin-free medium; growth is supported for approximately 90 min before the decline is observable. **(C)** Post-fusion recovery. HeLa cells are fixed at 1, 2, 3, and 4 h after PEG-mediated fusion. mCherry-tagged *E. coli* (red), DAPI-stained nuclei (cyan). Arrows mark extracellular bacteria, still present at 1 and 3 h and not detected at 4 h, establishing 4 h as the recovery interval used in the final protocol. Scale bar, 50 µm. **(D)** Timing matrix. Cells were seeded and incubated for 12 to 24 h prior to fusion. Fusion was then performed using one of eight polyethylene glycol (PEG) conditions: 10% or 20% PEG-3350, PEG-4000, PEG-6000, or PEG-8000 and fixed after additional 4 or 24 h. This resulted in a total of 32 experimental combinations. Optimal conditions identified included 12 h for fusion, 10% PEG-3350 (20% PEG-3350 was also effective; 10% was selected as the milder condition), and 4-hour incubation with gentamicin. The outcomes of this optimization are detailed in a matrix presented in Suppl. Fig. S4. Image created with BioRender.

##### Medium combinations

We evaluated various media types to determine optimal maintenance conditions for the HeLa cell line used which ensured the simultaneous viability of the bacteria and, ultimately, the endosymbiont of *A. deanei*. These media are usually supplemented with FBS (Fetal Bovine Serum) for nutritional support and with Penicillin-streptomycin (P/S) to prevent bacterial contamination. DMEM medium is a commonly used medium in mammalian cell culture and was selected for the experiments.

For cellular fusion and to perform experiments with the synthetic endosymbiont or to facilitate an eventual stable symbiosis, maintaining bacterial or endosymbiont viability in mammalian cell culture for multiple hours, or even days, is required. Once introduced, we aimed to preclude bacterial extracellular growth, i.e. bacteria in the cell culture medium. For this, we tested various combinations of these media, as shown in Fig. 3A and Suppl. Fig. S3, to determine which combination supports bacterial growth for a few hours and ultimately kills extracellular bacteria. Combinations 4 and 5 (DMEM+FBS+P/S+Gentamicin 50 µg mL^-1^ and 100 µg mL^-1^) appear to meet our requirements, as they support bacterial growth for 90 min, after which OD_600_ declines (Fig. 3B). Combination number four with 50 µg mL^-1^ was carried forward, on the basis of the concentration already selected as part of the bacterial parameter optimisation. The rationale for selecting this particular combination is that it is a commonly used composition for cultivating healthy mammalian cells. Consequently, adding gentamicin to the established medium appears to be the most appropriate approach to facilitate co-culture of HeLa cells and bacteria.

##### Timing of seeding, fusion and recovery

We fixed the medium combinations, and for recovery, tested four time intervals from 0 to 4 h post-fusion to determine the minimum time required to eliminate extracellular bacteria, as shown in Fig. 3C. With the medium and recovery interval fixed, we screened the remaining timing parameters together with PEG identity and concentration.

We identified the optimal PEG type and concentration for bacteria in the bacterial parameter optimisation part, however Ikada et al. (1995) reported less fusion between mammalian cells below 30% w/w PEG^29^. This threshold however was established for fusing two mammalian cells and not for the delivery of bacteria. We therefore aimed to use lower PEG concentrations while also exploring higher molecular weight PEGs. Consequently, we screened combinations of PEG types and concentrations, seeding-to-fusion intervals, and post-fusion recovery intervals (Fig. 3D; Suppl. Fig. S4). Cells were seeded and incubated for 12 or 24 h before fusion, with one of eight PEG conditions: 10 or 20% PEG-3350, -4000, -6000 or -8000, and fixed after further 4 or 24 h, resulting in 32 combinations in total (Fig. 3D). Ten conditions gave consistent delivery, as shown in Suppl. Fig. S4. Lower yield was observed for PEG-6000 and above, consistent with the higher viscosity of these solutions, which limited contact between the two membranes. A seeding-to-fusion interval of 12 h was more permissive than 24 h.

Two methods consistently proved effective: applying 10% or 20% PEG-3350 twelve hours after seeding, followed by a 4 h incubation with gentamicin post-fusion. The 4 h recovery period was deliberately chosen to exceed the 2 h duration required for gentamicin to suppress extracellular growth in the initial phase, ensuring that clearance is complete prior to imaging. Additional experiments involving HeLa cells post-fusion at various intervals over 4 h revealed extracellular growth of *E. coli* even after 2 h, leading to the choice of a 4 h recovery period instead of the previously tested 1.5 h.

Out of 32 conditions, 10 conditions showed positive delivery as shown in Suppl. Fig. 4. From these 10 conditions, 10% PEG-3350, 12 h after seeding and 4 h recovery was selected for all subsequent experiments. This condition was the least concentrated and lowest molecular weight PEG giving consistent delivery, and HeLa cells retained normal morphology and adherence throughout the procedure, whereas higher concentrations and molecular weights are associated with more variable outcomes. Also, delivery was more reproducible across replicate wells, and fewer visibly stressed or rounded cells.The final protocol involves mixing bacteria at OD_600_ 0.5 with a 3:1 ratio of 10% PEG-3350, then applying this mixture to HeLa cells 12 h after the seeding for 4 min at room temperature. The cells are then incubated in OptiMEM for 30 min at 37°C, followed by a 4 h incubation in DMEM containing 50 µg mL^-1^ gentamicin (Fig. 4). These conditions were maintained for all later experiments.

**Figure 4.**
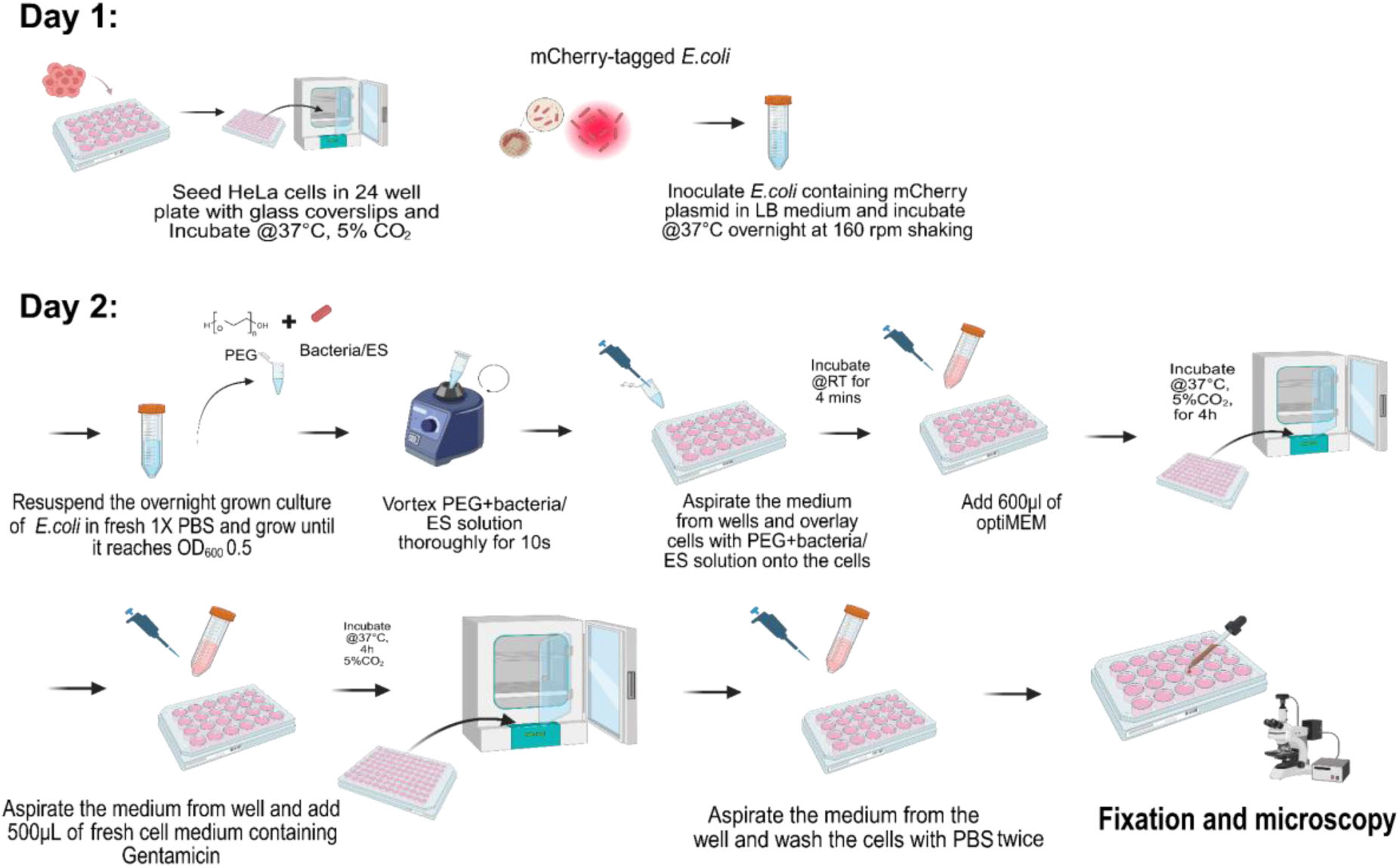
The resulting four-phase protocol. HeLa cells are seeded 12 h before the PEG fusion and *E. coli* is grown overnight; bacteria at OD_600_ 0.5 are mixed 3:1 with 10% PEG-3350 and applied for 4 min at room temperature, followed by addition of OptiMEM and incubation at 37°C, 5% CO_2_ for 30 min; extracellular bacteria are cleared in 50 µg mL^-1^ gentamicin for 4 h; cells are then fixed, DAPI stained and imaged. Image created with BioRender.

#### Final, optimised PEG-mediated *E. coli* - HeLa cells fusion protocol

The following describes the final, resulting protocol following the optimisation procedures described above and detailed in Suppl. Fig. S1-S4. The complete workflow is summarised in Fig. 4.

##### 1. Preparation of host cells and bacteria

Approximately 40,000 HeLa cells are seeded per well onto 24-well plates with glass-coverslips in a total volume of 500 µL of DMEM medium supplemented with 125 U mL^-1^ penicillin, 125 U mL^-1^ streptomycin and 10% FBS. The cells are then incubated at 37°C in a 5% CO_2_ atmosphere for approximately 12 h. On the same day, an *E. coli* culture is inoculated in LB medium and grown aerobically at 37°C, with shaking at 160 rpm

##### 2. Fusion of bacteria and HeLa cells

The overnight *E. coli* culture is pelleted at 6,000 *g* for 5 min and resuspended in PBS. OD_600_ is determined and adjusted to 0.5. A 400 µL 10% PEG 3350-bacteria mixture is prepared at a 3:1 ratio and vortexed for 10-15 s. The medium is aspirated from the HeLa cells, and then the cells covered with 400 µL of a PEG-bacteria mixture per well. The plate subsequently incubated at room temperature. After 4 min, 600 µL of Opti-MEM medium is added, and the mixture is incubated at 37°C in a 5% CO_2_ atmosphere for 30 min.

##### 3. Suppression of extracellular *E. coli* and recovery of the host

After 30 min of incubation, the medium is exchanged with DMEM supplemented with 125 U mL^-1^ penicillin, 125 U mL^-1^ streptomycin, 50 µg mL^-1^ gentamicin and 10% FBS. Next, the cells are incubated at 37°C in a 5% CO_2_ atmosphere for 4 h to recover. This step is performed to suppress the growth of extracellular *E. coli* cells. After 4 h of incubation, the medium is aspirated from the well and cells washed twice with 500 µL of DMEM supplemented with 125 U mL^-1^ penicillin, 125 U mL^-1^ of streptomycin and 10% FBS to get rid of residual extracellular bacteria.

##### 4. Fixation and imaging

After washing, the medium is aspirated from the well, and the cells are fixed with 4% paraformaldehyde for 10 min on ice, followed by 10 min at room temperature. Cells are stained with 250 µL of DAPI solution (1:1,000 dilution of a 1 mg mL^-1^ stock) for 10 min and washed once with 1X PBS to remove residual DAPI. Coverslips are embedded in Mowiol 4–88 supplemented with 15 mg mL^−1^ 4-diazabicyclo[2.2.2]octane and mounted on object slides. Samples are imaged on a confocal microscope system, Ti2 inverted microscope. mCherry, EGFP, and DAPI are visualised using excitation lasers of 561 nm, 488 nm, 408 nm, and emission filters of 570–1,000 nm, 500–550 nm, 417–477 nm, respectively. Image acquisition is performed using NIS-Elements. Analysis and processing (brightness and contrast) are adjusted using Omero Viewer (4.20.00).

### Implementation of the protocol

#### The optimised protocol delivers *E. coli* into HeLa cells

We first used *E. coli* as a generally prototyping platform because it is fast-growing and easy to work with, Applying the optimised protocol to mCherry-containing *E. coli* TOP10, bacteria were detected in association with HeLa cells by confocal microscopy after the gentamicin clearance step (Fig. 5A). To distinguish delivered bacteria from the bacteria adhering to the cell surface, we acquired z-stacks through the full depth of the host cells and generated orthogonal x-and y-projections in Fiji (Fig. 5B). Persistence of delivered bacteria was observed over 3-4 days. Delivery of bacteria was observed in HeLa cells across two independent experiments (n=2).

**Figure 5.**
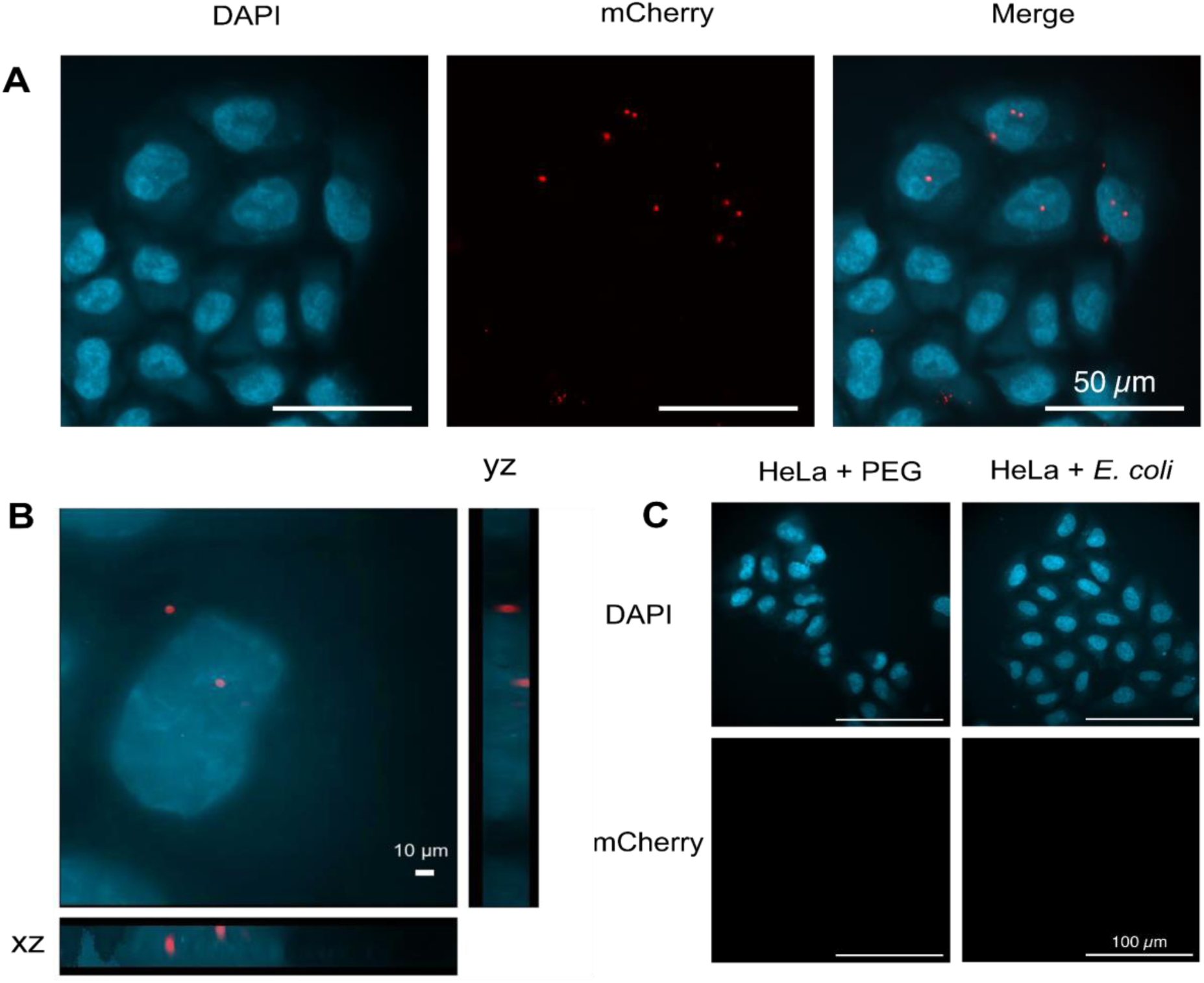
Optimised protocol delivers *E. coli* into HeLa cells. **(A)** Representative confocal images of HeLa cells 4 h after fusion with *E. coli* TOP10 expressing mCherry plasmid, DAPI (cyan, nuclei), mCherry (red, *E. coli),* and merge. Scale bar, 50 µm. **(B)** Orthogonal (xy, xz, yz) projections of a selected representative single HeLa cell, showing mCherry-positive *E. coli* signal confined to a specific region within the cell in all three planes. Scale bar, 10 µm. **(C)** Controls. HeLa cells treated with PEG alone (no bacteria), and HeLa cells incubated with mCherry-tagged *E. coli* without PEG. No intracellular mCherry signal was observed in either control. n=2 independent experiments.

#### Extension to the natural endosymbiont *Ca*. K. crithidii

Having established the protocol with *E. coli*, we asked whether it is readily applicable to a bacterium of direct evolutionary interest, *Ca.* K. crithidii; the obligate ß-proteobacterial endosymbiont of *A. deanei*. Isolation of *Ca*. K. crithidiii from *A. deanei* is laborious (see the materials and methods section), and yields material that is unstable outside its host cell. Because PEG-fusion requires no particular genetic modification of the bacterium, it is in principle applicable. For fusion of this endosymbiont with the HeLa cell, we used freshly isolated endosymbionts which were purified using density gradient centrifugation as discussed in the materials and methods section (Fig. 6A-B). Delivery of the endosymbiont was observed in HeLa cells across two independent experiments (Fig. 6C).

**Figure 6.**
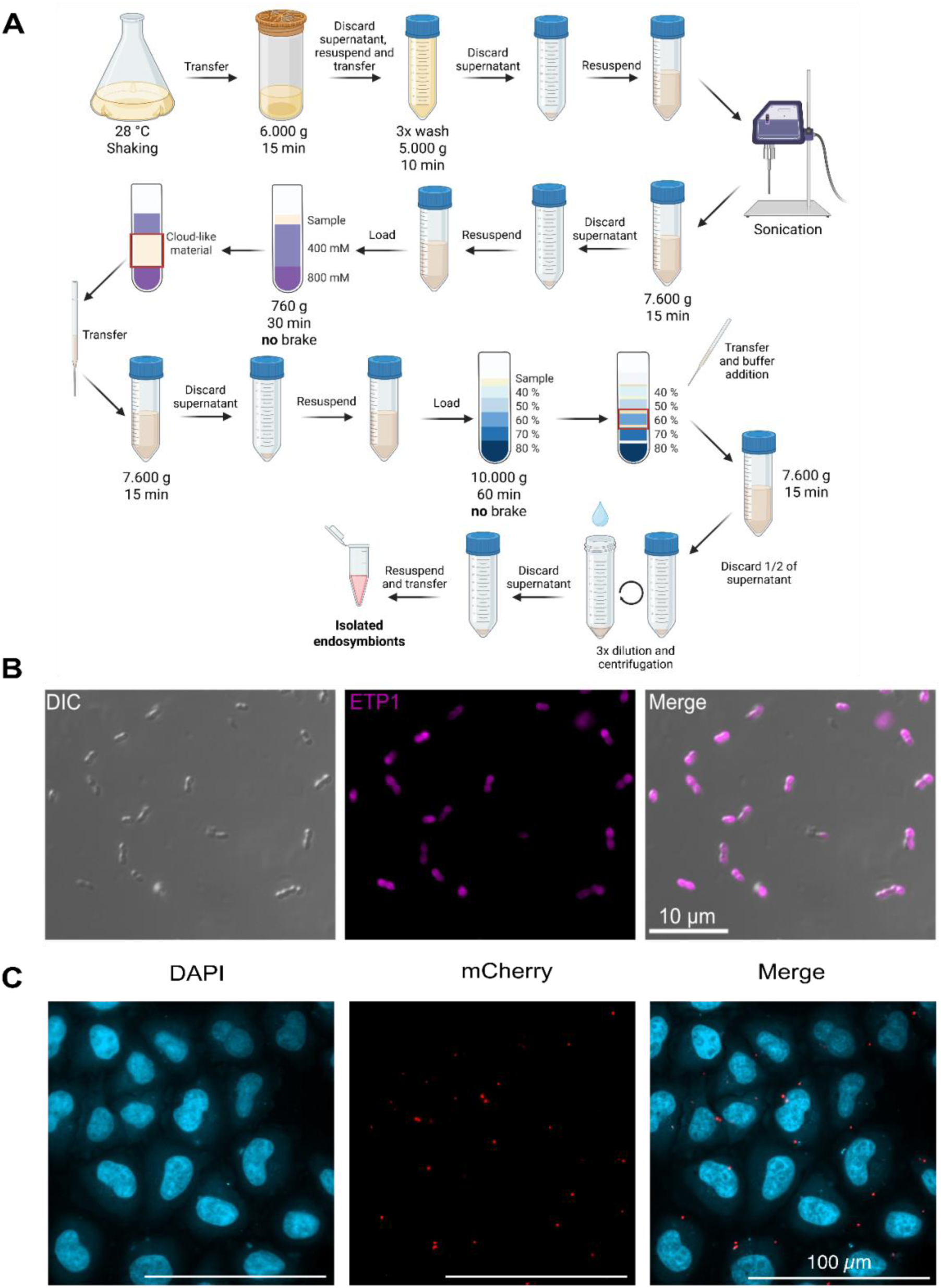
The applicability of the developed fusion protocol extends to the endosymbiont *Ca*. K. crithidii isolated from *A. deanei.* **(A)** Isolation workflow: *A. deanei* grown to late-log phase is harvested, lysed by sonication, and the endosymbiont fraction is purified through successive sucrose and Percoll PLUS gradients; freshly isolated material was used for PEG fusion. Created in BioRender. https://BioRender.com/s91nlns. (**B)** Micrograph of the isolated endosymbiont fraction. Endosymbionts identified by the typical peanut shape (DIC) and the mScarlet-tagged ETP1 marker, scale bar, 10 µm. **(C)** Confocal images of HeLa cells after the fusion with isolated *Ca*. K. crithidii, scale bar, 100 µm, n=2 independent experiments.

### Troubleshooting

Table 1 lists the drawbacks most frequently encountered during development of this protocol, together with the likely causes and the solutions we adopted.

**Table 1:**
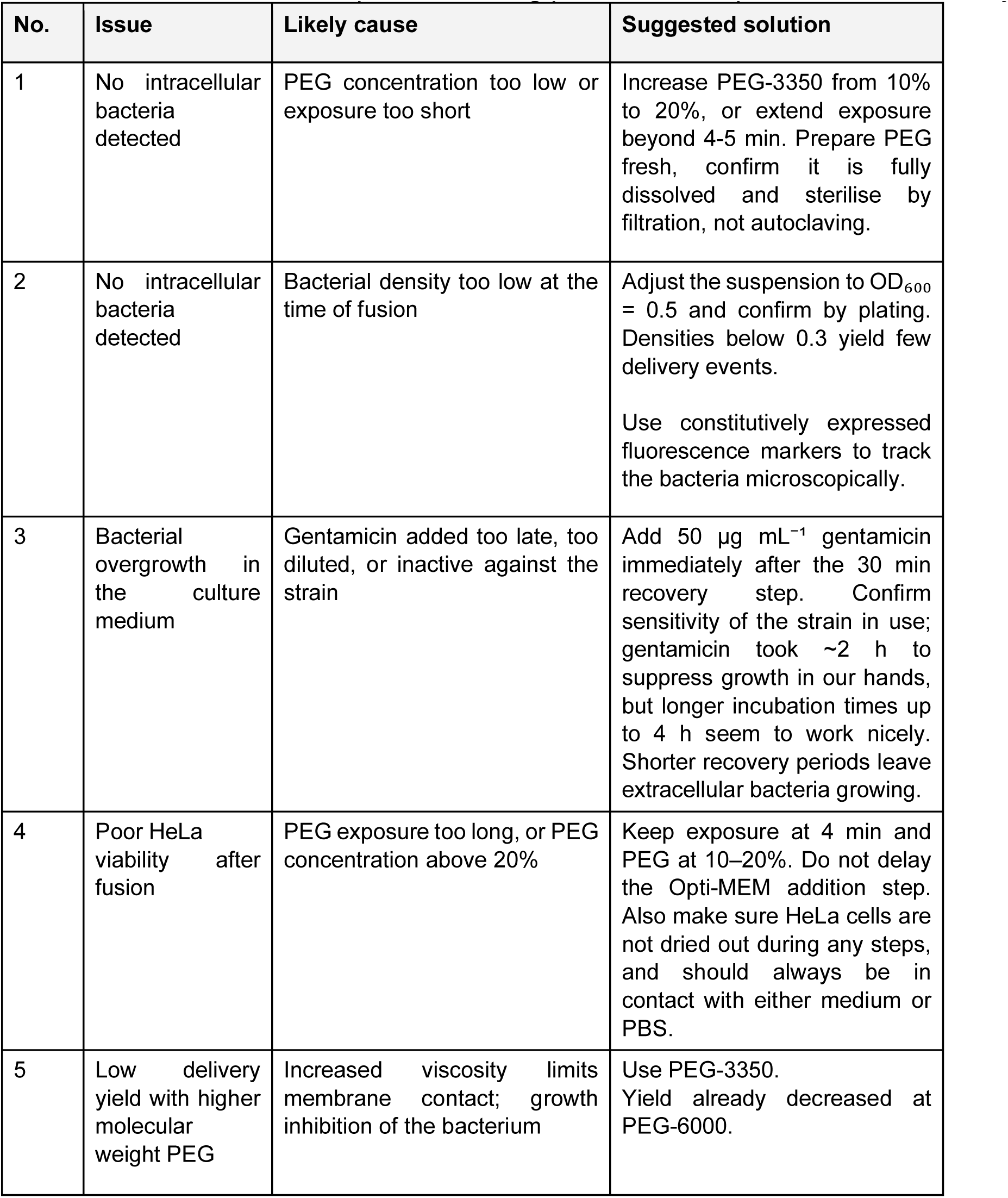
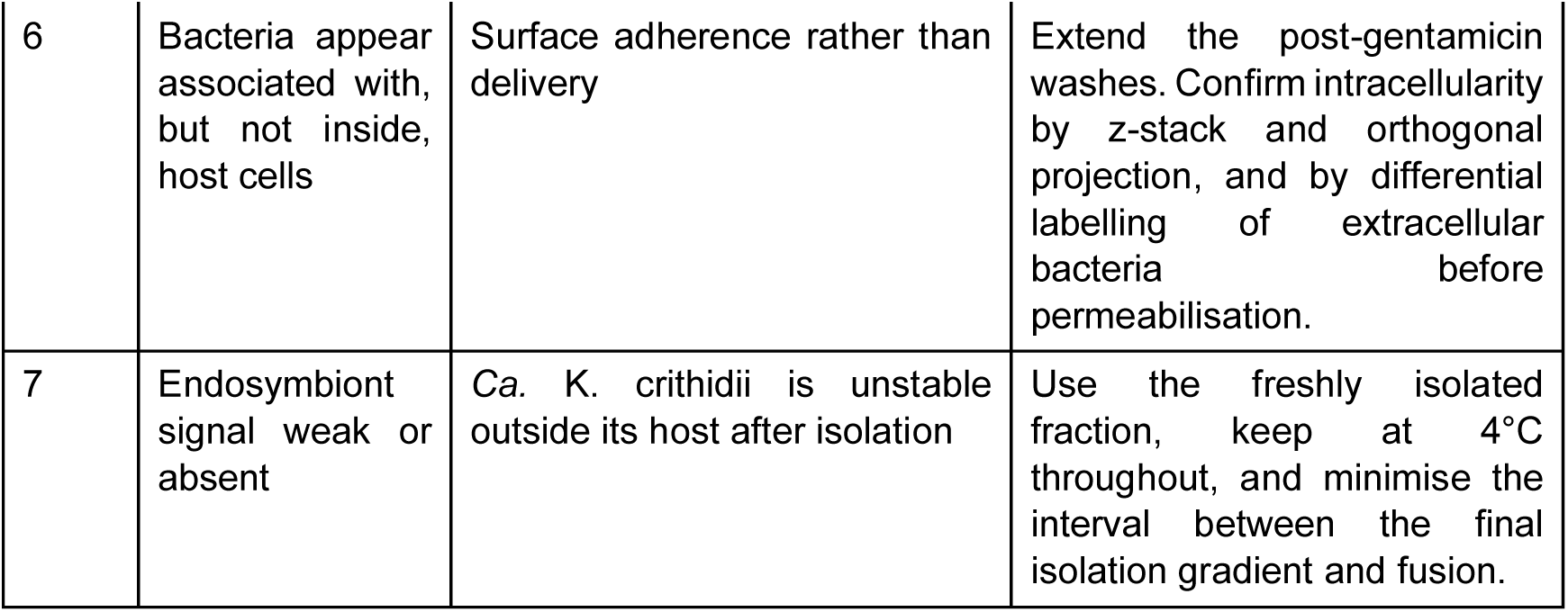
Troubleshooting guide for PEG-mediated delivery of bacteria into mammalian cells. Issues are listed in the order in which they arise during the protocol. Causes and solutions are based on the experience during protocol development in our laboratory.

### Remarks and limitations of the protocol

The compartment in which the delivered bacteria reside has not been determined yet, z-stack reconstruction places them inside the host cell but cannot distinguish cytosol from a host-derived vacuole, and this distinction matters for any downstream endosymbiosis analysis or follow up experiment. Consequently, all claims in this study are limited to a localisation within the host cell.

The viability of bacteria following delivery has not been directly evaluated. The persistence of the mCherry fluorescence signal over three to four days suggests potential survival but does not confirm bacterial viability, as fluorescent proteins can remain detectable in non-metabolically active bacteria. No assertion is made regarding whether the delivered bacteria are alive, dividing, or metabolically active.

HeLa cells retained normal morphology and adherence post-procedure, but these observations are qualitative. The protocol is a bulk process with a stochastic outcome, so the population is heterogeneous. Some cells receive no bacteria while others receive several, and we have not established what determines this heterogeneity. As we have not examined the bacteria or endosymbiont persistence beyond a few days, no claim is made about maintenance over host cell division. The viability of the extracellular bacteria is not assessed. OD_600_ measures the turbidity which also includes the non-viable cells and debris, so the data establish that extracellular bacteria cease to grow rather than they are removed. Recovery of colony-forming units after the clearance step would distinguish the two.

We are interested whether a specific cell cycle stage of the host cell favours bacterial or endosymbiont introduction. The two bacterial species tested differ in aspects that may influence the generalizability of the findings. *E. coli* was employed for the protocol optimisation due to its availability, ease to culture. The size of *Ca*. K. crithidii is similar to *E. coli*.

Lastly, the experiments were conducted with a single host cell line; HeLa cells were selected for their nutrient rich cytosol, helpful for the growth of bacteria and flattened morphology and large cytoplasmic volume, which facilitate fusion and imaging. The applicability of the protocol to smaller, rounder, less adherent, or immunologically competent cell types remains to be evaluated.

## Discussion and perspectives

We describe here a polyethylene glycol-based method that results in the introduction of bacteria into mammalian cells. Our goal was to identify conditions in which fusion occurs while both the bacteria and host cells remain viable. We observed that 10% PEG-3350 facilitated efficient delivery, whereas concentrations of 30% or higher halted bacterial growth. Timing is crucial: gentamicin takes about 2 h to eliminate extracellular bacteria, establishing a minimum recovery period, while extended PEG exposure improves delivery at the expense of bacterial viability. Each parameter is limited by the balance between fusion needs and cellular tolerance; the resulting general protocol optimally intersects these constraints rather than maximizing one. The main advantage over existing approaches is that neither partner needs to be engineered. Microinjection reliably introduces bacteria but targets individual cells and requires specialised equipment^8,9^. Methods such as engineered invasion and listeriolysin-mediated phagosomal escape can be scaled up to populations but depend on the endosymbiont being culturable and genetically tractable^7,10^. This is a significant constraint, not just a technical inconvenience; the genome reduction that indicates an endosymbiont’s advanced integration stage toward becoming an organelle also hinders its independent cultivation and subsequent engineering. Organisms further along the integration pathway, which are most informative about the process, are largely inaccessible to the most effective methods. PEG fusion overcomes this limitation; the bacterium is a passive participant, making tractability irrelevant. Additionally, this is a bulk process performed while isolated endosymbionts remain viable, which is crucial for materials that cannot be prepared in advance or in large quantities.

A second potential advantage, based on the underlying mechanism, still needs to be tested. In invasion pathways, bacteria are enclosed in a host vacuole and must escape afterwards; this explains why listeriolysin O is co-expressed with invasin in engineered systems^7^, and why phagolysosomal degradation remains the main barrier to endosymbiosis in mammalian cells^5^. PEG acts by directly fusing adjacent bilayers, possibly allowing direct incorporation of the bacteria into the host cell cytosol without a vacuole, thereby bypassing the escape challenge. However, PEG treatment may also promote uptake into a membrane-bound compartment via a route similar to macropinocytosis, in which the barrier is simply delayed rather than removed^32^. Co-staining with a plasma membrane marker and an endolysosomal pathway marker could indicate whether a host membrane surrounds the bacterium. Stimulated emission depletion (STED) microscopy and correlative light and electron microscopy (CLEM) would together eventually establish the identity and ultrastructure of any eventual bacteria-surrounding membrane.

Bacterial and host viability after the delivery also requires direct assessment.. Recovering colony-forming units from lysed host cells or inducing a reporter after delivery would directly demonstrate viability. Live-cell imaging over comparable intervals would show whether the delivered bacteria divide and would indicate whether the host tolerates their presence throughout its own division cycle. Host viability and proliferation after PEG exposure should be quantified alongside delivery efficiency rather than inferred from morphology, particularly since some PEG derivatives have been reported to affect HeLa cell growth^30^. The outcomes of the protocol vary across the population. A hypothesis worth exploring, the host cell cycle stage might influence permissiveness, as membrane composition, surface area, and cortical organization change throughout the cycle. Differentiating between possibilities is crucial both practically and mechanistically, since a protocol that consistently delivers a single bacterium per cell would be more effective for studying integration than one with variable numbers. Delivery is the first of several requirements for establishing an endosymbiotic relationship, and this paper focuses solely on that aspect. Persistence through host cell division, coordinating replication with the host, achieving metabolic alignment via transporters, and ultimately transferring genes to the host nucleus remain unaddressed^5,33^. The historical context of PEG-based associations illustrates the scope of this challenge: HeLa-tobacco hybrid lines, although proliferative, retained only fragments of the plant genome^22^. Advances have been made through alternative approaches; Gäbelein et al. (2022)^10^ demonstrated endosymbiotic growth of *E. coli* within mammalian cells^10^ and Courneyer et al. (2022)^34^ achieved photosynthetic functionality via engineered endosymbiosis, both methods involving significant bacterial modification^34^. Our contribution emphasises not greater persistence but broader accessibility, a delivery method applicable to organisms that cannot be genetically engineered and effective at the population level.

For a generalised application of the protocol, as for instance here illustrated, one primary goal might be to characterise host integration of the endosymbiont, i.e. *Ca*. K. crithidii in greater detail. Unlike *E. coli*, this organism has spent its evolutionary history within a eukaryotic cytosol and has lost the genes for functions that its host supplies. Whether that history helps or hinders its survival in an unrelated host is unknown, and either outcome would be informative. If its persistence is similar to that of *E. coli*, it would indicate that adaptation to an intracellular environment is transferable between hosts, implicating general features of the cytosol rather than anything specific to *A. deanei.* Poorer persistence would indicate that the dependence is specific, and would help identify what the trypanosomatid host provides that HeLa cells lack.

More broadly, because the method requires no engineering of either partner, it should be transferable readily to other intractable bacteria, and to other mammalian cell types, providing a general starting point for reconstructing host-endosymbiont integration.

## Materials and Methods

### Bacterial cell culture

Two *E. coli* strains were used: T7 express *E. coli* carrying plasmid pET28a-sfGFP for IPTG-inducible sfGFP expression, and OneShot TOP10 (Invitrogen cat. no. C4040-03, competency: >10^−9^ CFU μg^−1^), constitutively expressingmCherry (Addgene, pmr101A-cPr-mCherry, cat. No. 177207). The T7 Express strain was used for all optimisation experiments; whereas the TOP10 strain was used for fusion, where its constitutively expressed mCherry complements the mScarlet-tagged endosymbiont. Cultures were inoculated from a glycerol stock into 5 mL of Luria-Bertani broth (Roth, cat. no. 6673.4) containing 50 µg mL^-1^ ampicillin for T7 express *E. coli* along with 1.5 mM IPTG and for oneshot TOP10 *E. coli* cells, 5 mL of Luria-Bertani broth (Roth, cat. no. 6673.4) containing 50 µg mL^-1^ kanamycin and cultures were incubated overnight at 37°C with shaking at 160 rpm.

### Growth curve

An overnight-grown culture was inoculated into a fresh LB or DMEM-based medium. The OD_600_ was either adjusted for the OD optimization step, or it was maintained at 0.5 throughout the experiment. For every condition, 2 mL of mixture was prepared. 150 µL aliquots were dispensed in triplicate into a 96-well flat-bottom plate, and OD_600_ was recorded at 0, 30, 60, 90, 120, 150 and 180 min using a plate reader CLARIOstar (BMG LABTECH, Ortenberg, Germany). Plates were held at 37℃ with shaking at 160 rpm between the readings. All the readings were blanked against the cell-free medium of the corresponding composition.

### Gentamicin screen

Gentamicin (Sigma, Cat. no. G1397) was added at 10, 25, 50 or 100 µg mL^-1^ to bacterial suspensions at OD_600_ 0.15, 0.2, 0.3 and 0.5, resulting in 16 conditions (Suppl. Fig. S1).

### PEG screen

PEG-3350,-4000, -6000 and -8000 were tested at 10, 20, 30, 40, 50% (w/v) in bacterial cultures at OD_600_ 0.5, with each molecular weight with its own no-PEG control, resulting in 20 conditions in total (Suppl. Fig. S2). PEG solutions were prepared in PBS, sterile-filtered, and used fresh. Readings in PEG-containing medium were blanked against the corresponding PEG-only control.

### Media screen

Five cumulative DMEM-based compositions were compared at a starting OD_600_ of 0.5: (1) DMEM; (2) DMEM + 10 % FBS; (3) DMEM +10% FBS + 125 U mL^-1^ Penicillin and streptomycin; (4) DMEM + 10% FBS + 125 U mL^-1^ Penicillin and streptomycin + 50 µg mL^-1^ gentamicin; (5) DMEM + 10% FBS + 125 U mL^-1^ Penicillin and streptomycin + 100 µg mL^-1^ gentamicin. Media were prewarmed to 37°C before incubation (Fig. 3A-B, Suppl. Fig. S3). Mean 士 SD, n=3

### Mammalian cell culture

HeLa (DSMZ,ACC 57) cells were cultivated in a humidified atmosphere at 37℃, 5% CO_2_ in Dulbecco’s modified Eagle medium (DMEM, PANBiotech, Aidenbach, Germany, cat. no. P04-03550), supplemented with 125 U mL^-1^ penicillin, 125 U mL^-1^ streptomycin (PANBiotech, cat. no.P06-07100), and 10% fetal bovine serum (FBS, PAN Biotech, cat. no. P30-3602).

### Cultivation of *Angomonas deanei* and endosymbiont extraction

Endosymbionts were isolated essentially as described previously^15^. In brief, an *A. deanei* strain expressing the endosymbiont marker mScarlet-ETP1^32^ was grown in Brain Heart Infusion (BHI, Sigma-Aldrich) containing 10 µg mL^-1^ hemin (Sigma-Aldrich) at 28°C with shaking. To isolate a sufficient number of endosymbionts at least 300 mL of a late-log culture were used for extraction. All isolation steps were carried out at 4°C. Cells were harvested by centrifugation for 15 min at 6,000 *g* followed by three washing steps with 50 mL 1x PBS and 10 min of centrifugation at 5,000 *g* each. The cell pellet was resuspended in 15 mL of buffer A (10 mM HEPES, 0.5 mM EDTA and 20 mM KCl, pH 7.4) containing 150 mM sucrose and a protease inhibitor cocktail (cOmplete™, Mini, EDTA-free Protease Inhibitor Cocktail, Roche; in addition, 1 mM PMSF, 10 µM E64 and 100 µM Na-tosyl-L-lysine can be added). Cells were lysed by sonication with 10 cycles, an amplitude of 80 %, 0.8 cycles, and 18 pulses per cycle using a Hielscher UP50H Compact Lab Homogenizer equipped with an MS3 sonotrode. Sufficient cell lysis was confirmed microscopically. The cell lysate was centrifuged at 15 min at 7,600 *g* and the supernatant discarded. The pellet was resuspended in 8 mL buffer A containing 250 mM sucrose. The suspension was split and 4 mL each loaded on top of two discontinuous sucrose gradients prepared with 5 mL buffer A containing 800 mM sucrose and 10 mL buffer A containing 400 mM sucrose in 30 mL glass tubes (Corex). The gradients were centrifuged at 760 *g* for 30 min without brake in a Beckman JS-21 centrifuge with a swinging-bucket rotor JS-13.1. After centrifugation the entire cloud-like material in the upper half of the gradient was transferred to a 50 mL tube and centrifuged at 7,600 *g* for 15 min. The pellet was resuspended in 2 mL buffer A containing 250 mM sucrose. The suspension was split and 1 mL each loaded on top of two discontinuous Percoll® PLUS (Cytiva) gradients of 2 mL each 80 %, 70 %, 60 %, 50 % and 40 % Percoll in buffer A with a final concentration of 250 mM sucrose prepared in 15 mL glass tubes (Corex). Gradients were centrifuged at 10,000 *g* for 1 h without brake in a Beckman JS-21 centrifuge with a swinging-bucket rotor JS-13.1. The endosymbionts are contained in the fraction that is visible at/between the 60 % - 70 % Percoll bands. This fraction was transferred to a 50 mL reaction tube and an equal volume of buffer A containing 250 mM sucrose was added. The suspension was centrifuged at 7,600 *g* for 15 min. Afterwards, half of the supernatant was removed and the removed volume was substituted with the same volume buffer A containing 250 mM sucrose. This step was repeated three times until a visible pellet formed. Most of the supernatant was removed carefully and the pellet was resuspended in the remaining supernatant (∼200 - 500 µl). The suspension was analysed microscopically to verify that endosymbionts were successfully isolated and OD_600_ was measured. Images of isolated endosymbionts were acquired with an Axio Imager A.2 and processed with the Zen Blue software v2.5 (both from Zeiss). For fusion experiments, freshly extracted endosymbionts were used.

### Post-fusion recovery imaging

PEG-mediated fusion was performed as described below and replicates were fixed after 1, 2, 3 and 4 h of recovery in gentamicin-containing medium. Extracellular bacteria were identified as mCherry positive rod. Three fields were scored per time-point from 2 independent experiments (Fig. 3C).

### Timing Matrix

HeLa cells were seeded and incubated (as mentioned before) for 12 to 24 h before fusion. The PEG fusion was performed with each one of the eight conditions (10 or 20% PEG-3350, -4000, -6000 or -8000), and fixed after a further 4 or 24 h incubation period, resulting in 32 combinations (Fig. 3D; Suppl. Fig. S4). Delivery of bacteria in mammalian cells was scored using confocal microscopy as a presence of intracellular mCherry or sfGFP signal. Conditions with no detectable intracellular signal under these criteria were scored as negative. Three fields were scored per time-point from 2 independent experiments.

### Quantification of delivery

Delivered bacteria were observed using confocal z-stacks acquired throughout the depth of the host cell. A bacterium was counted as intracellular when sfGFP/mCherry signal was resolved in optical sections interior to host cell in both orthogonal projections. Four cells were scored per conditions across 2 independent experiments.

### Statistics

No inferential statistical tests were conducted. All comparisons are descriptive. For the growth assays (Figs. 2B, 2C, 3B, Suppl. Fig. S1-S3), each screen was performed as three independent experiments; the figures show data from one representative experiment, with values reported as mean ± SD of three technical replicates measured in parallel wells of that experiment. Error bars therefore reflect within-experiment (well-to-well) variability rather than variability between independent experiments. The same overall pattern was observed in all three independent experiments. In imaging experiments, n indicates the number of fields or cells scored, as noted in each legend. Post-fusion recovery (Fig. 3C) was assessed from three fields per timepoint across two independent experiments. Timing matrix conditions (Fig. S4) and delivery of *Ca.* K. crithidii (Fig. 6) were assessed across two independent experiments. Results are thus reported as observed outcomes rather than quantified efficiencies. Growth curve means and SDs were calculated in R version 4.6.1 using dplyr version 1.2.1, and figures were produced with ggplot2 version 4.0.3.

## Acknowledgements

We are grateful to S. Kuschel and R. Schönle (Heinrich-Heine-Universität Düsseldorf, Germany) for valuable experimental assistance.

## Supplementary information

**Suppl. Fig. S1.**
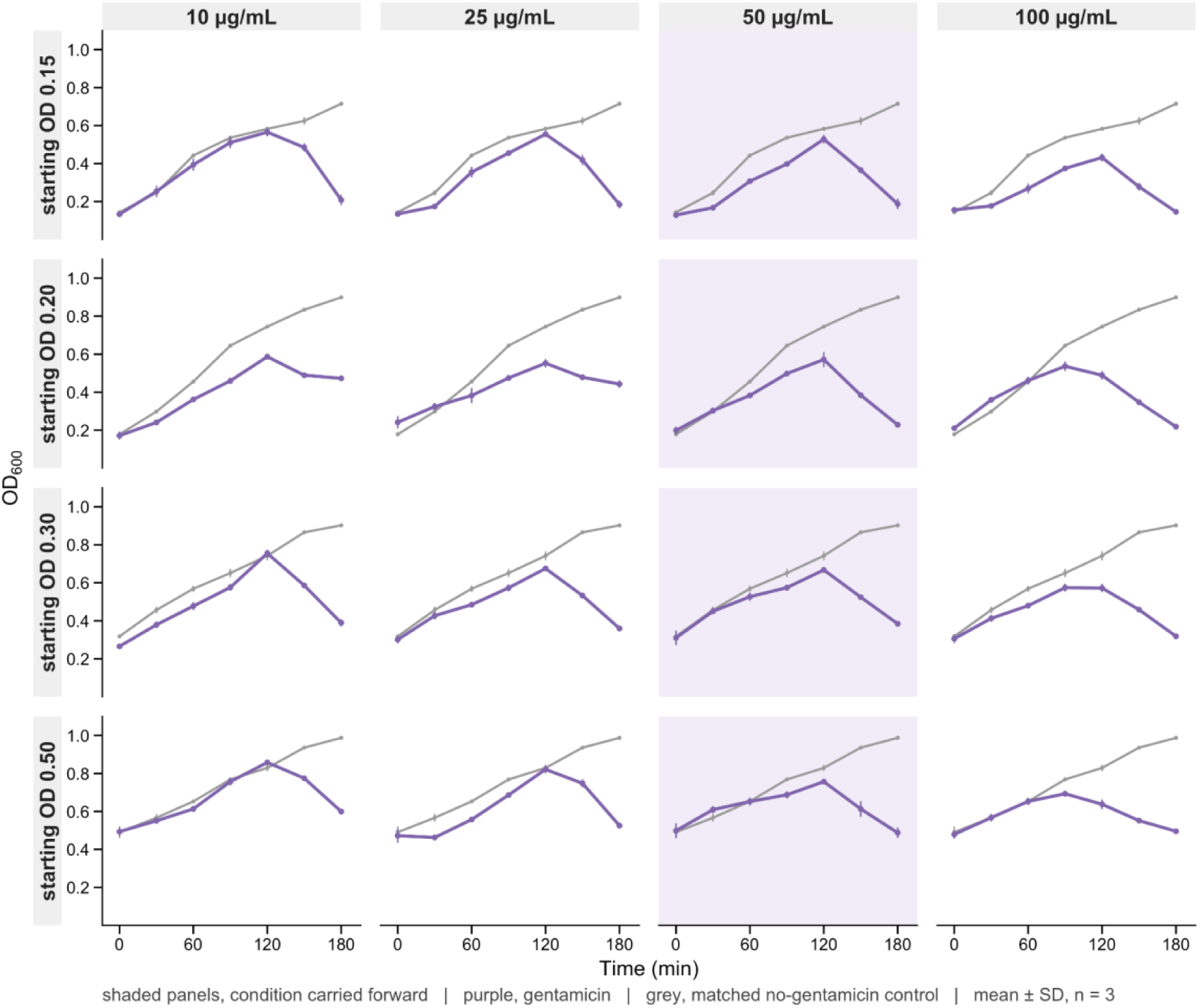
Complete gentamicin vs bacterial density matrix: Growth of *E. coli* at gentamicin concentrations of 10, 25, 50, and 100 µg mL^-1^ (columns) and OD_600_ of 0.15, 0.2, 0.3, and 0.5 (rows). Each panel shows the gentamicin (purple) with the corresponding no-gentamicin (grey) growth curve, repeated across four panels of each row. All gentamicin concentrations suppressed growth, with OD_600_ peaking at 120 min and then declining thereafter, whereas controls continue to increase throughout. 50 µg mL^-1^ gentamicin (shaded column) was selected as the lowest concentration that provided consistent suppression across the full density range and carried forward for subsequent experiments. Mean 士 SD, n=3.

**Suppl. Fig. S2.**
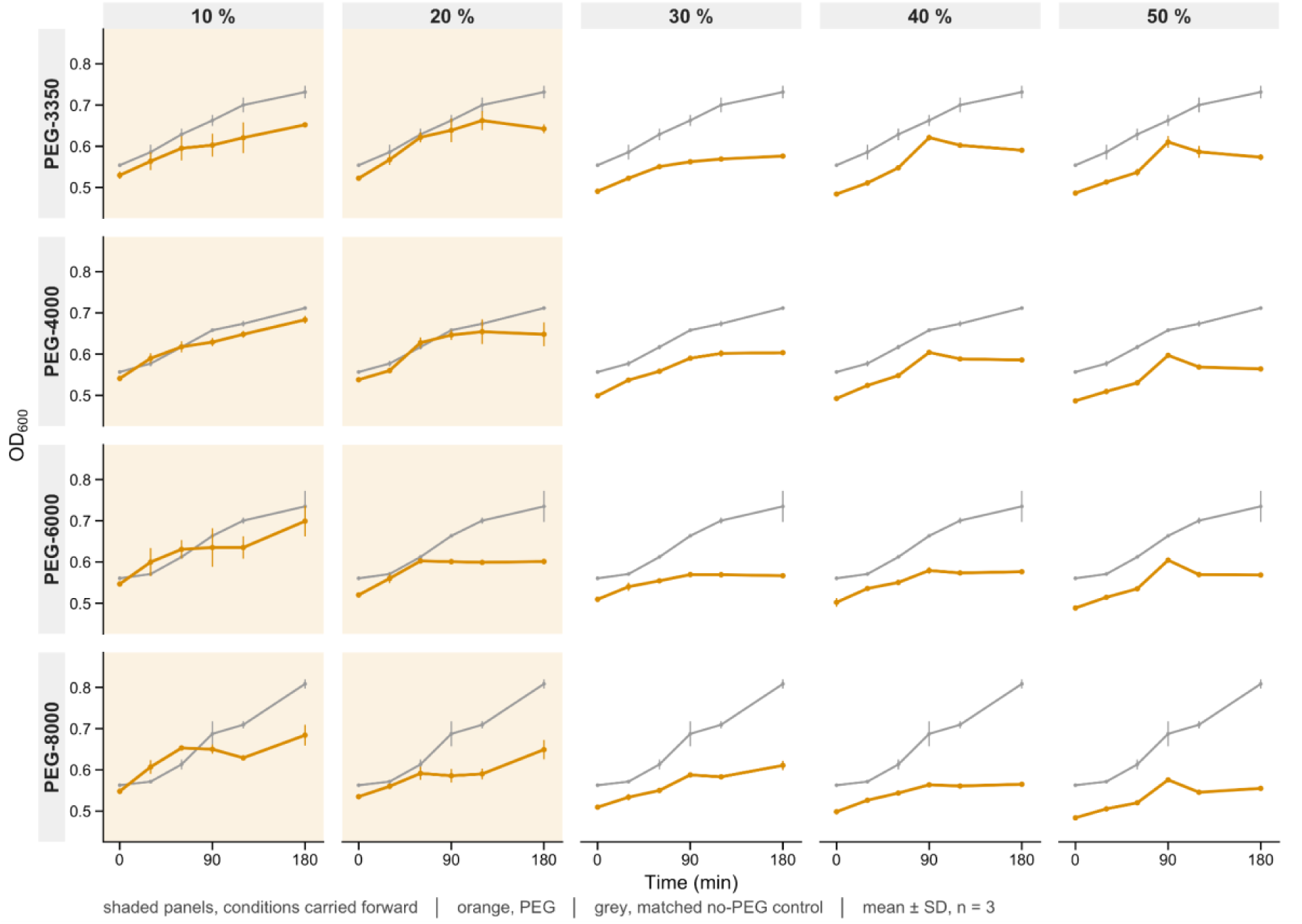
Complete PEG molecular weight vs. concentration screen matrix. Growth of *E. coli* at OD_600_ 0.5 with PEG-3350, PEG-4000, PEG-6000, and PEG-8000 at 10, 20, 30, 40, and 50% recorded over 180 min. Each panel shows the PEG condition (orange) with the corresponding no-PEG (grey) control; the same control trace is repeated across the five panels in each row. Growth was reduced relative to control at every concentration tested, and the reduction increased with concentration. Molecular weight had little effect. The shaded panel indicates the ten conditions carried forward into host and fusion optimisation experiments. Mean 士 SD, n=3.

**Suppl. Fig. S3.**
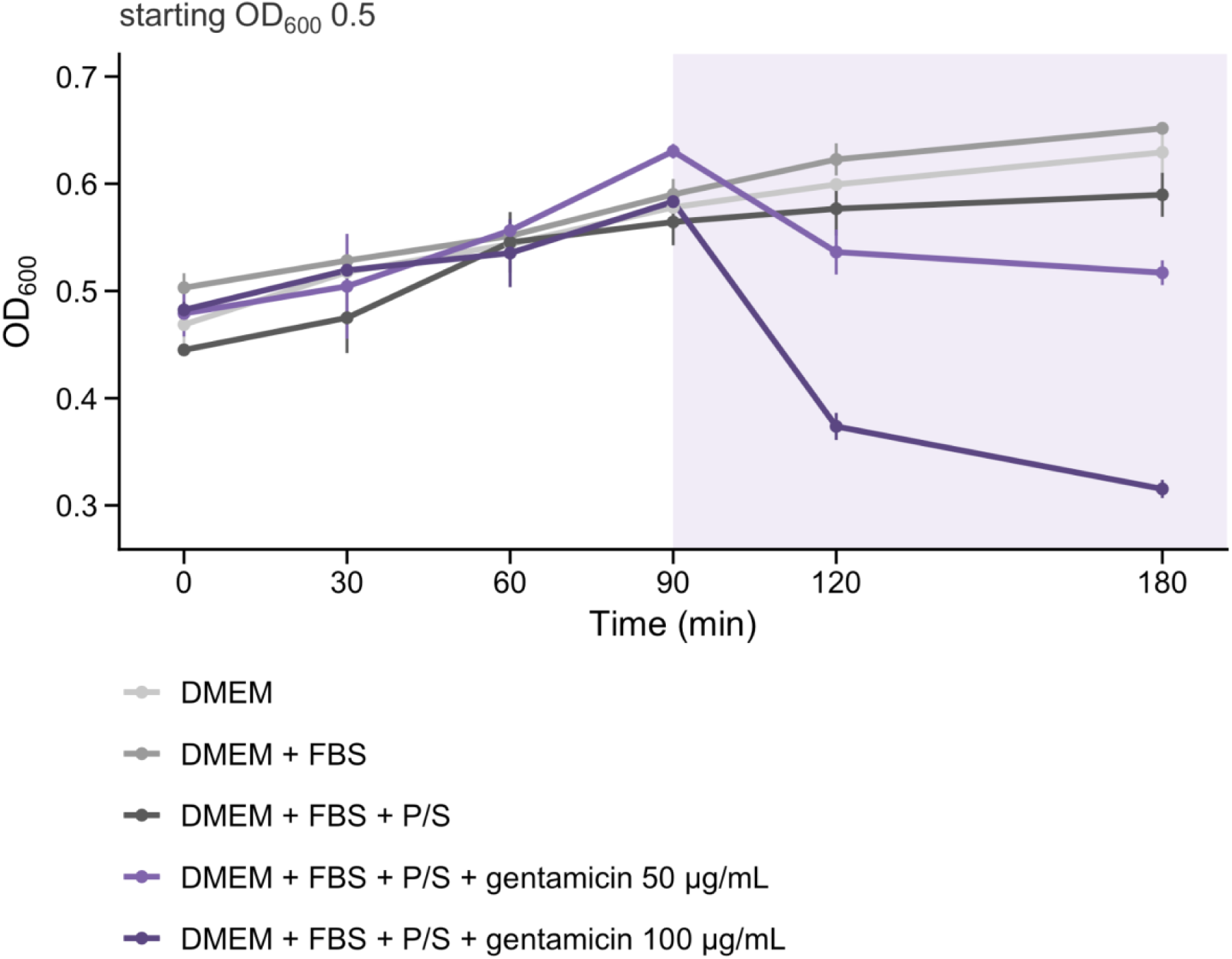
Complete medium panel. Growth of *E. coli* at a starting density of OD_600_ 0.5 in five DMEM based combinations: (1) DMEM; (2) DMEM+FBS; (3) DMEM+FBS+Penicillin-Streptomycin; (4) DMEM+FBS+Penicillin-Streptomycin+50 µg mL^-1^ gentamicin; (5) DMEM+FBS+Penicillin-Streptomycin+100 µg mL^-1^ gentamicin. Growth was unaffected by the addition of FBS. Penicillin-streptomycin led to a slower growth compared to the first two combinations; only the gentamicin containing combinations showed a reduction, with OD_600_ peaking at 60 min and declining thereafter. Combination 4 was carried out in the final protocol, on the basis of a gentamicin concentration selected in Fig. 2B. Mean 士 SD, n=3.

**Suppl. Fig. S4.**
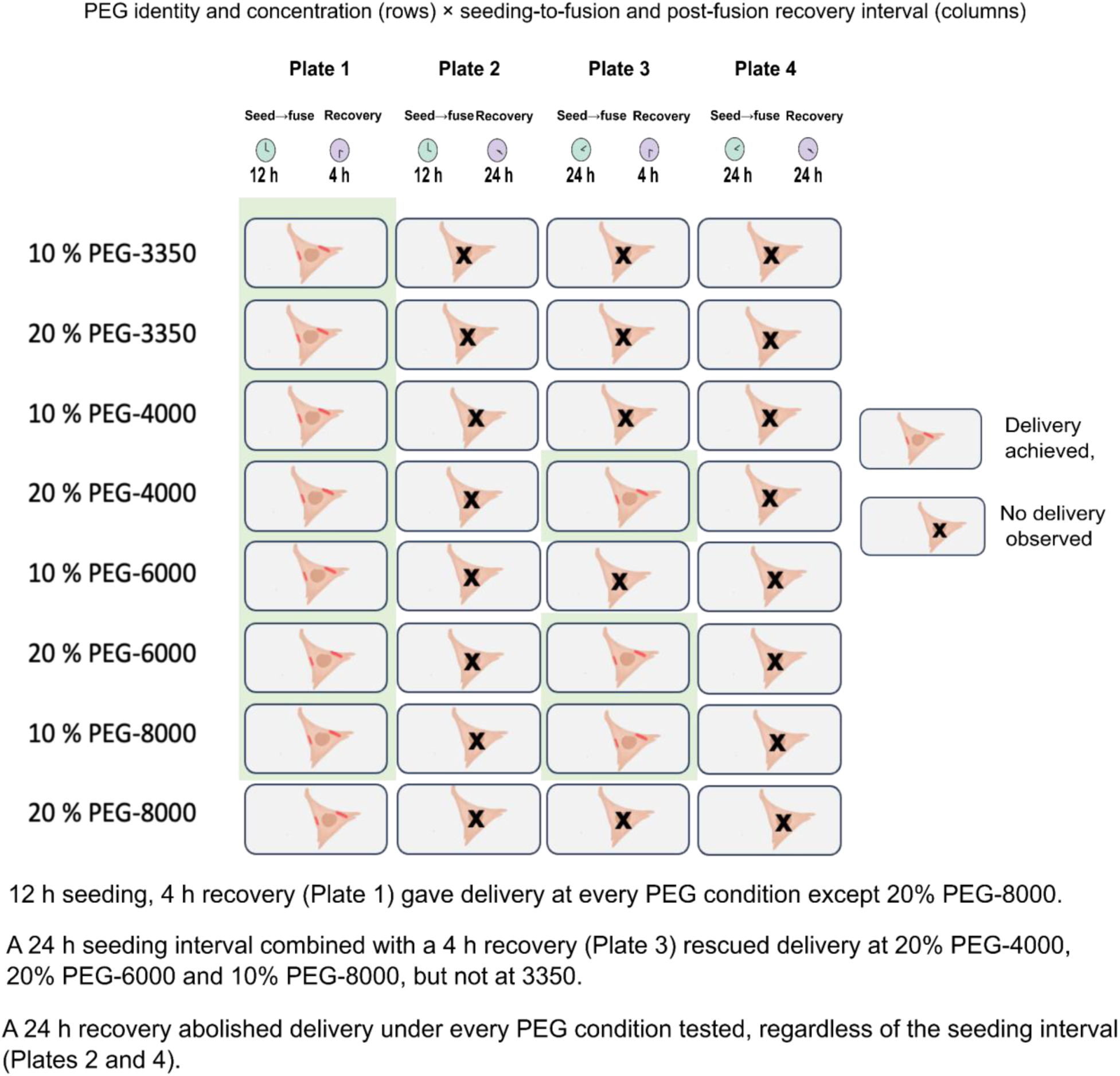
Complete timing matrix. All 32 tested combinations with PEG type and concentration, seeding-to-fusion interval and post-fusion recovery interval, with outcome scored for each. Out of these 32 conditions, 10 were scored positive. Out of these ten, 10% PEG-3350 was selected for the final protocol. Image created with BioRender.

**Supplementary Table S1.** Strains, plasmids and reagents used in this study.

Bacterial strains
| Strain | Source | Used for |
| --- | --- | --- |
| *E. coli* T7 Express | NEB (cat. No. C2566H) | Optimisation (density, gentamicin and PEG screens) |
| *E. coli* One Shot TOP10 | Invitrogen (cat. no. C4040-03) | PEG-mediated fusion into HeLa cells |

Trypanosomatid host and endosymbiont
| Organism | Strain/line | Source | Used for |
| --- | --- | --- | --- |
| *Angomonas deanei* | mScarlet-ETP1 expressing strain based on ATCC PRA-265 from the American Type Culture Collection | Generated in Morales et al; 2016 | Source of mScarlet-ETP1-marked *Ca. K. crithidii* |

Mammalian cell line
| Cell line | Description | Source | Used for |
| --- | --- | --- | --- |
| HeLa | Human cervical epithelial adenocarcinoma | DSMZ, ACC 57 | PEG-mediated fusion, all optimisation and imaging experiments |

Plasmids
| Strain | Genotype/plasmid | Source | Used for |
| --- | --- | --- | --- |
| pET28a-sfGFP | IPTG-inducible sfGFP, kanamycin resistance | Addgene, cat.no. 85492 | Optimisation |
| pmr101A-cPr-mCherry | Constitutive mCherry, ampicillin resistance | Addgene, cat. no. 177207 | Final PEG fusion protocol |

Reagents
| Reagent | Supplier | Catalogue no. | Used for |
| --- | --- | --- | --- |
| Luria-Bertani broth | Roth | 6673.4 | Bacterial culture |
| Gentamicin | Sigma-Aldrich | G1397 | Suppression of extracellular bacterial growth |
| Ampicillin | Sigma-Aldrich | A9518 | T7 Express selection |
| Kanamycin | Sigma-Aldrich | K4000-25G | TOP10 selection |
| IPTG | Thermo Fisher scientific | R0392 | sfGFP induction |
| Polyethylene glycol (PEG) | Sigma | 3350: [c] 4000: [cat. no.]<br>6000: [cat. no.] 8000:<br>[cat. no.] | Fusion reagent |
| DMEM | PAN Biotech | P04-03550 | HeLa culture and fusion medium |
| Fetal bovine serum (FBS) | PAN Biotech | P30-3602 | HeLa culture medium |
| Penicillin–streptomycin | PAN Biotech | P06-07100 | HeLa culture medium |
| Opti-MEM | Gibco | 22600-134 | Post-fusion dilution |
| Paraformaldehyde, 4% in PBS | VWR | J61899 | Cell fixation |
| DAPI | Sigma-Aldrich | D9542-5MG | Nuclear counterstain |
| Mowiol 4-88 | Carl Roth | 0718 | Mounting medium |
| DABCO | Carl Roth | 0713 | mounting medium |
| Percoll | Cytiva | 17544501 | Density-gradient isolation of *Ca. K. crithidii* |

